# Altered meningeal immunity contributing to the autism-like behavior of BTBR *T*^+^ *Itpr3^tf^*/J mice

**DOI:** 10.1101/2022.01.29.478292

**Authors:** Mohammad Nizam Uddin, Kevin Manley, David A. Lawrence

## Abstract

Autism spectrum disorder (ASD) is a complicated neurodevelopmental disorder, which is categorized by deficiency of social contact and communication, and stereotyped forms of performance. Meningeal immunity conditions the immune reflection and immune defense in the meningeal area involving meningeal lymphatic organization, glymphatic structure, immune cells, and cytokines. The development of meningeal immunity dysfunction might be the leading cause for many neural diseases including ASD. The inbred mouse strain BTBRT+*Itpr3tf*/J (BTBR) shows multiple ASD-like behavioral phenotypes, thus making this strain a widely used animal model for ASD. In our previous study, we reported an altered peripheral immune profile in BTBR mice. Herein, we are investigating immunological and neural interactions associated with the aberrant behavior of BTBR mice. BTBR mice have an increased level of immune cell deposition in the meninges along with a higher level of CD4^+^ T cells expressing CD25 and of B and myeloid cells expressing more MHCII than C57BL/6 (B6) mice, which have normal behaviors. BTBR mice also have higher levels of autoantibodies to dsDNA, Aquaporin-4, NMDAR1, Pentraxin/SAP and Caspr2 than B6 mice, which may affect neural functions. Interestingly, the T regulatory (Treg) cell population and their function was significantly reduced in the meninges and brain draining lymph nodes, which may explain the increased level of activated B and T cells in the meninges of BTBR mice. A low level of Treg cells, less IL-10 production by Treg, and activated T and B cells in meninges together with higher autoantibody levels might contribute to the development of autism-like behavior through neuroinflammation, which is known to be increased in BTBR mice.

**Highlights:**

1. BTBR mice have higher level of immune cell deposition in the meninges compared to C57BL/6 (B6) mice.
2. Meningeal T cells and B cells of BTBR mice express a higher level of CD25 and MHCII, respectively, than those of B6 mice.
3. BTBR mice have a higher level of serum autoantibodies to dsDNA and brain antigens (Aquaporin-4, NMDAR1, Pentraxin/SAP and Caspr2) than B6 mice.
4. T regulatory (Treg) cell population was reduced in the meninges and brain draining lymph nodes of BTBR mice with lower cytokine production of IL-10.
5. Fewer Treg cells and more activated meningeal T and B cells together with higher autoantibody levels might contribute to the development of the autism-like behavior of BTBR mice.

## Introduction

Autism spectrum disorder (ASD) is characterized by the existence of numerous different indicators such as verbal impairment, deficiency in social interaction, and monotonous pattern of behavioral. ASD is a lifelong neurodevelopmental condition with a strong involvement of dysfunctional or altered immune system. Currently, in USA, 1 out of 68 children aged 8 years have been diagnosed with ASD [1–3]. The pathogenesis/progression of ASD involves a dysfunctional immune system including activation of both innate and adaptive immune cells. Dysregulated T cell subsets such as Th1, Th2, Th17 and Treg have been associated with ASD [4–7].

Meningeal immunity is important for neuronal homeostasis and for neuronal activity through the neuro-modulatory cytokines affecting neuronal signaling, animal behavior, senses and thought [8–10]. The meninges neighboring the brain are occupied by a diversity of immune cell types, which not only offer immune observation but also affect brain function [11]. Recently, it has been reported that meningeal lymphatic system can regulate neuronal lymphatic drainage and neuroinflammation [12]. Meningeal inflammation caused by various agents can influence neurological disorders and plays a key role in governing immunity in the central nervous system [13]. Recent investigation of COVID-19 has been suggesting neural malfunctions further connecting neuroimmune and vascular functions [14–16].

The meningeal lymphatic system is connected to the peripheral space and could sample and drain T cells, B cells, myeloid cells and CSF contents directly from the deep cervical lymph nodes [17,18]. Meningeal lymphatic system ensures metabolic homeostasis between the parenchyma and peripheral tissue; and play a role in regulating immune surveillance and immune responses in central nervous system [19,20]. Several brain functions such as spatial learning, short-term memory sensory responses and hippocampal neurogenesis are controlled by meningeal T cells through the secretion of cytokines. [21–24]. IL-4 and IL-13 secreted by CD4^+^T cells can promote astrocytes to express brain derived neurotrophic factor (BDNF) [21, 25], and IL-4 stimulates microglia to produce BDNF, IGF-1, and TGF-β to affect neuronal activity [26,27]. Additionally, meningeal macrophages are reactive to the condition of the surrounding situation and can control the immune responses by their plasticity of anti- or pro-inflammatory phenotypes [28]. Meningeal dendritic cells (DCs) can sense and carry any antigens (Ags) to peripheral T cells [29]. In addition, meningeal DCs might also induce peripheral tolerance by restraining T follicular helper (Tfh) and T follicular regulatory (Tfr) cell differentiation [30]. Moreover, impairment of meningeal lymphatics could cause weakened drainage of brain Ags, accrual of metabolic wastes and encourage immune cells to enter the brain region, unsettling the neuronal connections and leading to irregular behaviors [31–35].

T regulatory (Treg) cells are defined as a CD4^+^CD25^+^ T cell population expressing transcription factor forkhead box P3 (FoxP3) whose deficiency is linked to the development of severe autoimmunity [36–38]. Both soluble factors like cytokines and direct contact by cell-surface molecules between cells could possibly function as suppression molecules in Treg cell–mediated immune regulation [39]. Treg cells can prevent neuroinflammation through the regulation of Th17 cell effector functions by limiting access of Th17 cells to antigen presenting cells (APCs) and suppression of Th17 [40].

The BTBR mouse model has been increasingly used to study the underlying mechanisms for the development of autism [41–44], which shows numerous behavioral and immune aberrations detected in children with autism [45]. BTBR mice showed lack of social communication including increased level of monotonous self-grooming and nominal vocalization in social interaction [41,42]. Altered immune structure of Th1, Th2, Th17, and T regulatory cells along with cytokine, chemokine and transcriptional signaling was observed in the BTBR mice [4, 46].

Previously we reported higher levels of serum IgG and anti-brain antibodies (Abs), higher expression of some cytokines in the periphery and brain of BTBR mice [43] and maternal environment importantly maternal anti-brain autoantibodies (autoAbs) influence the development of ASD-like behavior [44]. An increase of Tfh cells and antibody producing plasma cells in BTBR mice was also reported [47]. Herein, we measure the immune profile of meningeal immune cells and serum autoAb levels.

## Results

### Immune cell profile in the meninges

Meningeal immunity can affect brain homeostasis and brain function by releasing neuromodulatory cytokines, controlling neuronal signaling, behavior, senses, and thought. An increasing number of studies are focusing on meningeal immunity as its dysfunction contributes to neurological diseases or neurodegeneration. In autism research, meningeal immunity is a poorly studied aspect of neuroimmune interactions. In our previous study, we observed altered immune profiles in peripheral blood and lymphoid organs. Herein, meningeal immunity in the BTBR autism mouse model is assayed. To investigate the immunoprofiles of meninges in B6 and BTBR mice, we analyzed the CD45^+^ total immune cells, CD3^+^ T cells, CD19^+^ B cells and CD11b^+^ myeloid cells by flow cytometry. The frequency of CD45+ cells in BTBR mice meninges was higher compared to B6 mice (Fig:1A), and the statistical analysis showed the deposition of immune cells in BTBR meninges was significantly higher than B6 mice (Fig:1D, P=0.005). When we further investigated the types of immune cells in meninges, we found that the frequency of CD3^+^ T cells and CD19^+^ B cells were significantly higher in BTBR meninges compared to B6 (Fig:1 B-D, P=0.0013 & P=0.0003); whereas the frequency of CD11b^+^ myeloid cells was not different in between B6 and BTBR mice. The higher deposition of total immune cells and higher frequency of T cell and B cell populations may shape the immunoprofile activities of the meninges in BTBR mice.

**Figure 1.**
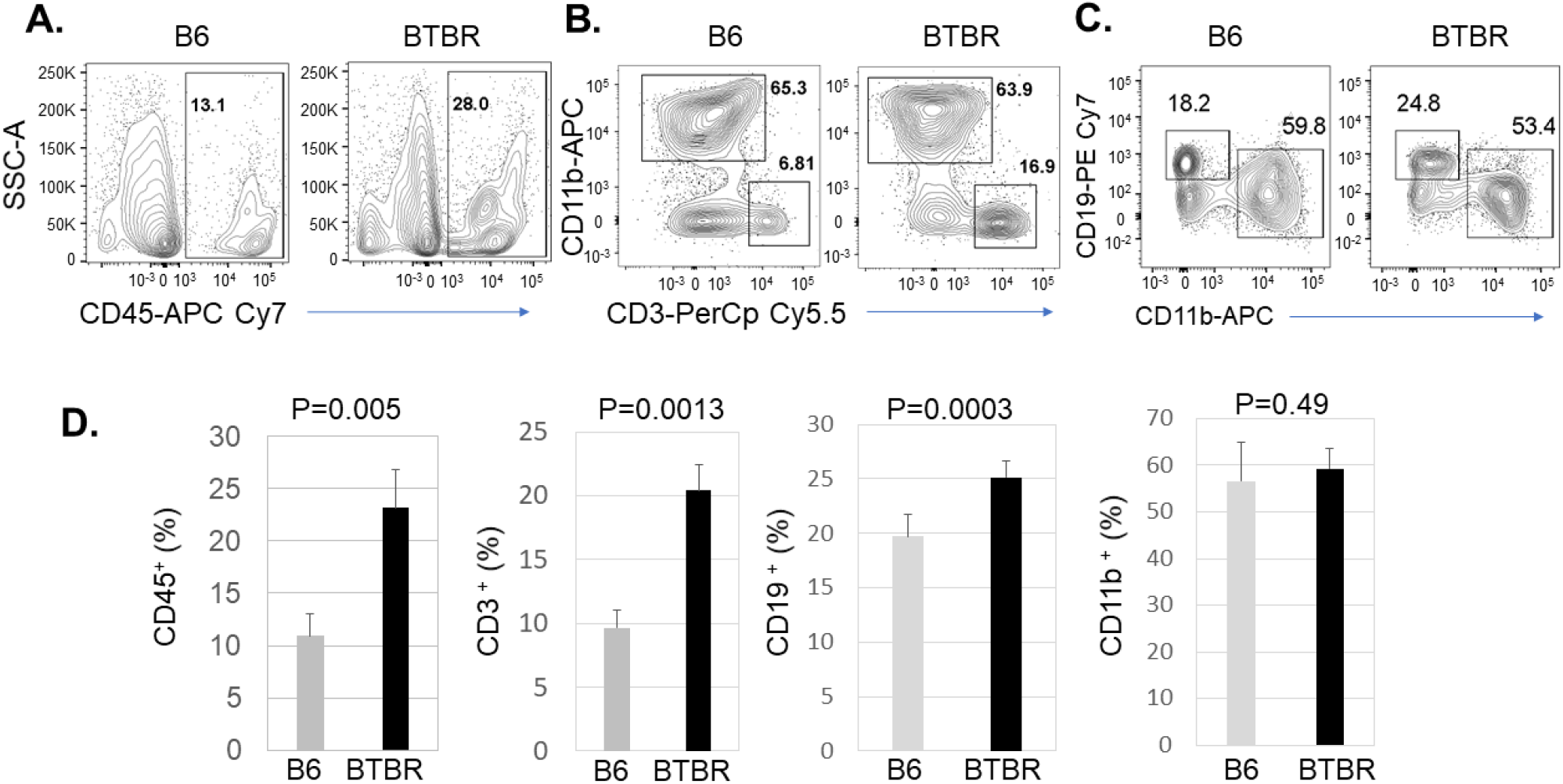
BTBR mice have significantly higher deposition of immune cells in meninges than B6 mice. Representative flow cytometric analysis of CD45^+^ cells (A), CD3^+^ T cells (B), CD19^+^ B cells and CD11b^+^ myeloid cells (C) populations in the meninges of B6 and BTBR mice. Frequency of CD45^+^ immune cells, CD3^+^ T cells, CD19^+^ B cells and CD11b^+^ myeloid cells (D) in the meninges of B6 and BTBR mice. Data are representative of three or four independent experiments with 4-6 pairs of mice in each experiment. The *p* values were determined by unpaired two-tailed Student’s t test, and *p*<0.05 is considered as significantly different. Error bar indicates mean ± SEM.

### Activation status of meningeal immune cells

To assess the function or activation conditions of the T cell and B cell populations that were increased in the BTBR mice, we evaluated the expression of CD25 on T cells and MHCII expression on B cells and myeloid cells. The expression of CD25 (Fig: 2A) significantly higher (P<0.0001) in BTBR mice indicating more active T cells in the meninges or possibly more Treg cells, which were assayed later. The expression level of MHCII also higher on B cells (Fig: 2B, P=0.0006) and CD11b^+^ myeloid cells (Fig: 2C, P<0.0001) of BTBR mice compared with B6 mice suggesting greater action and higher Ag presenting activity of B and myeloid cells in the meninges of BTBR mice compared to B6 mice.

**Figure 2.**
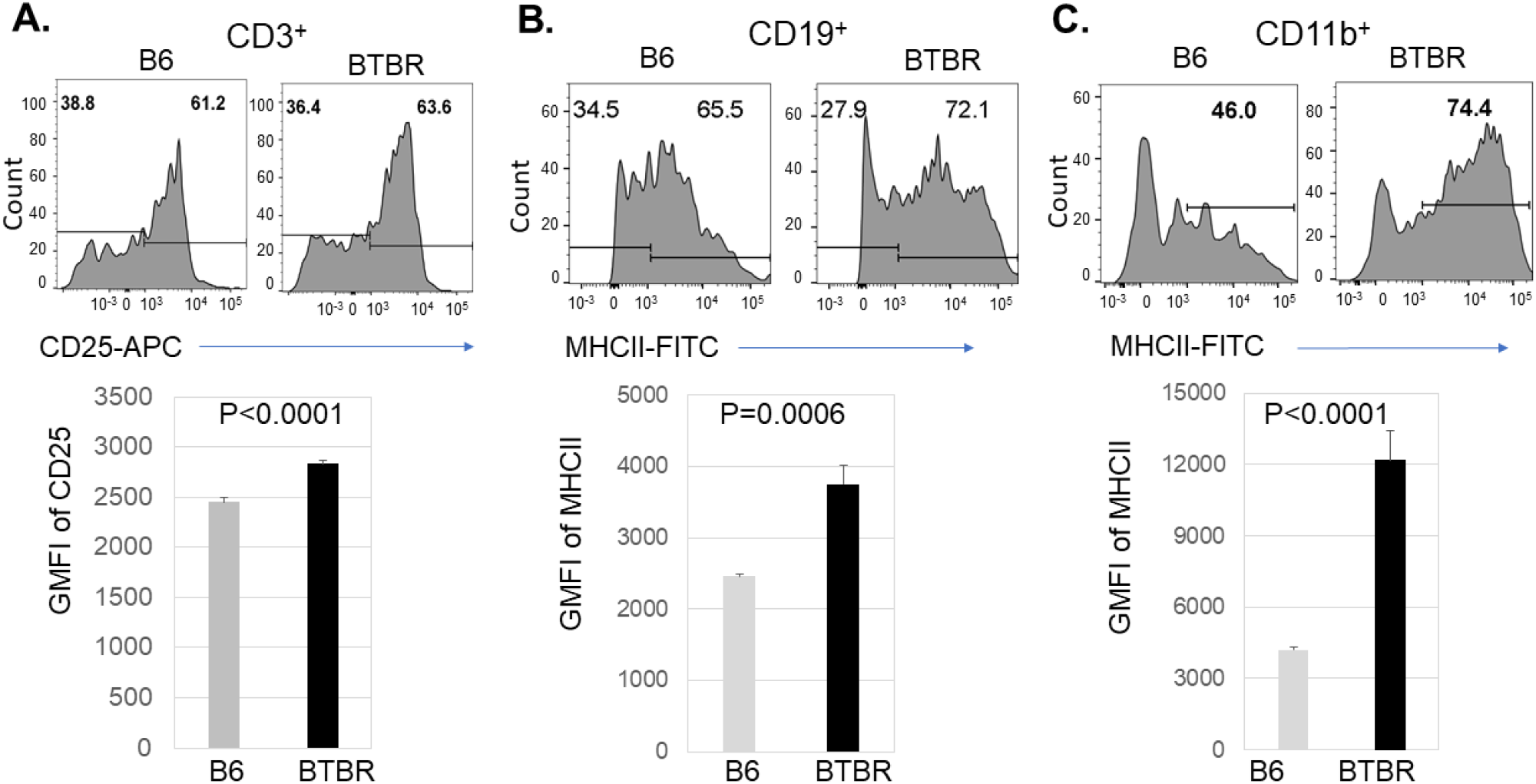
Increased expression of CD25 and MHCII on meningeal CD45^+^ cells of BTBR mice. Representative flow cytometric analysis of CD25 expression on CD3^+^ T cells (A), MHCII expression on CD19^+^ B cells (B) and CD11b^+^ myeloid cells (C) in the meninges of B6 and BTBR mice. Data are representative of three or four independent experiments with 4-6 pairs of mice in each experiment. The *p* values were determined by unpaired two-tailed Student’s t test, and *p*<0.05 is considered as significantly different. Error bar indicates mean ± SEM.

### Immune cell frequency in the cervical lymph nodes

Cervical lymph nodes are the main draining lymph nodes of meningeal lymphatic system. The meningeal lymphatic vessels oversee draining immune cells, small molecules, and additional fluid from the central nervous system into the cervical lymph nodes. Considering the high connection with the meningeal lymphatic system, we also investigated the immune profile of cervical lymph nodes. Like the meningeal system, we assayed the frequency of CD3^+^ T cell, CD19^+^ B cell and CD11b^+^ myeloid cell populations in cervical lymph nodes (Fig: 3A & B). The frequency of CD3^+^ T cells were significantly higher in the cervical lymph of BTBR mice compared to B6 mice (Fig: 3C, p=0.002). A similar profile was observed in BTBR meninges; whereas the frequency of CD19^+^ B cells in cervical lymph nodes (Fig: 3C, P=0.0001) was reduced in BTBR mice compared to B6 mice, which is opposite of meninges where it was increased. Like meninges, the frequency of CD11b^+^ myeloid cells (Fig: 3C, p=0.76) were not different in the cervical lymph nodes of B6 and BTBR mice.

**Figure 3:**
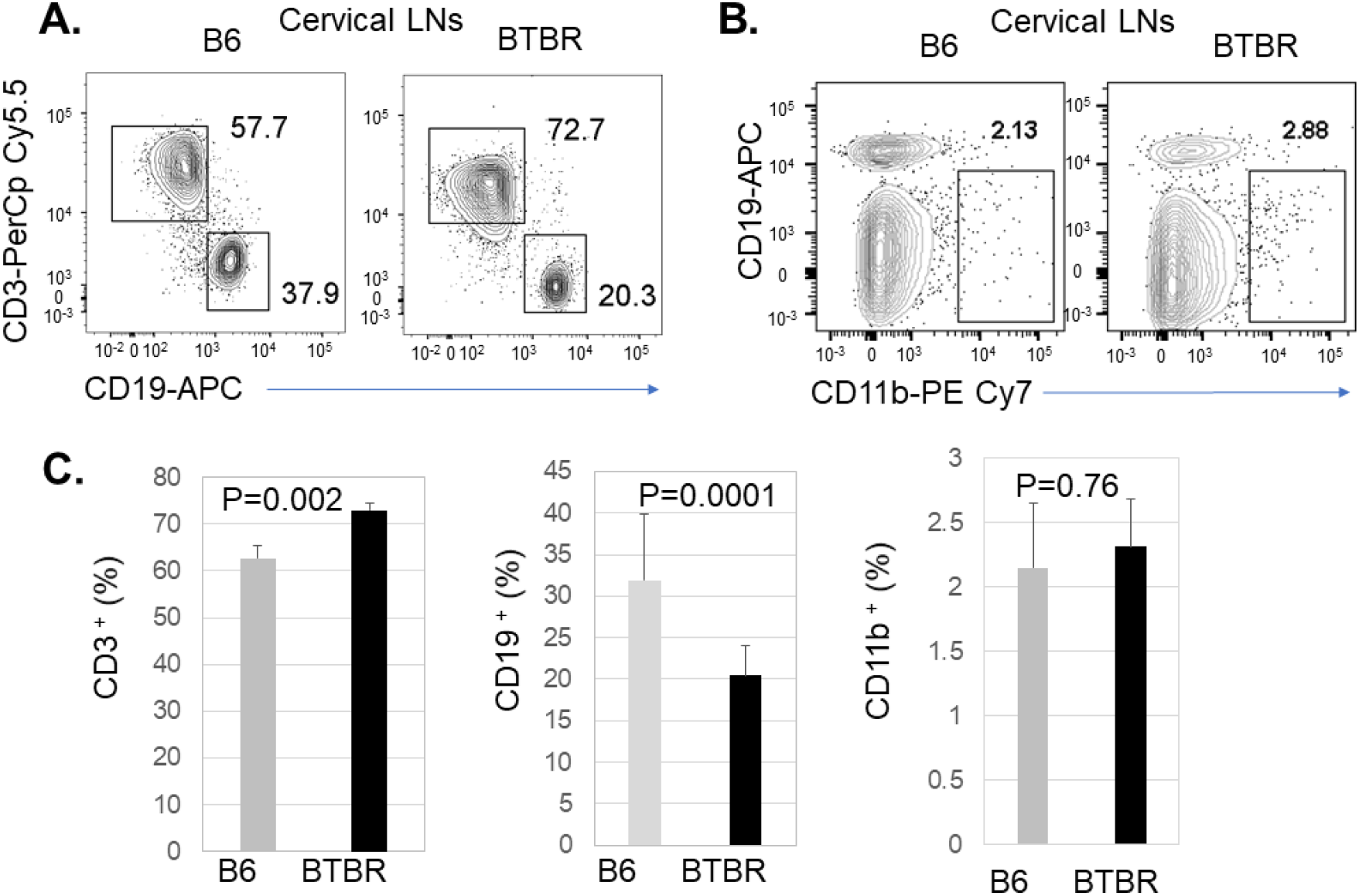
BTBR mice have significantly higher frequency of T cells and lower frequency of B cells in the cervical lymph nodes than B6 mice. Representative flow cytometric analysis and frequency of CD3^+^ T cells (A), CD19^+^ B cells (A), and CD11b^+^ (B) myeloid cells in the cervical lymph nodes of B6 and BTBR mice. Data are representative of three or four independent experiments with 4-6 pairs of mice in each experiment. The *p* values were determined by unpaired two-tailed Student’s t test, and *p*<0.05 is considered as significantly different. Error bar indicates mean ± SEM.

### Plasma cell and B cells in the cervical lymph nodes

Like the reduced frequency of B cells in the cervical lymph nodes, a similar trend was observed in BTBR spleens [48]. While B cell frequency was reduced, plasma cells were significantly higher in BTBR spleens [47] suggesting the lower frequency of B cells in the cervical lymph nodes may also be due to increased differentiation to plasma cells. Therefore, the frequency of plasma cells in cervical lymph nodes was investigated. The analysis the plasma cells population was accomplished by quantification of CD138^+^ cells in the CD3^-^CD19^-^ population by flow cytometry. The frequency of CD138^+^ plasma cells was significantly higher in the cervical lymph nodes of BTBR mice (Fig: 4A, p=0.01) compared to B6 mice. The expression of CD40 on B cells is essential for the generation of germinal centers, isotype switching, and stable antibody secretion. To investigate B cell activity, CD40^+^ B cells and the expression level of CD40 on B cells was assessed. Both, the frequency of CD40^+^ B cells and the expression levels of CD40 (Fig: 4B, P<0.0001) were higher on BTBR B cell compared to those of B6 mice.

**Figure 4:**
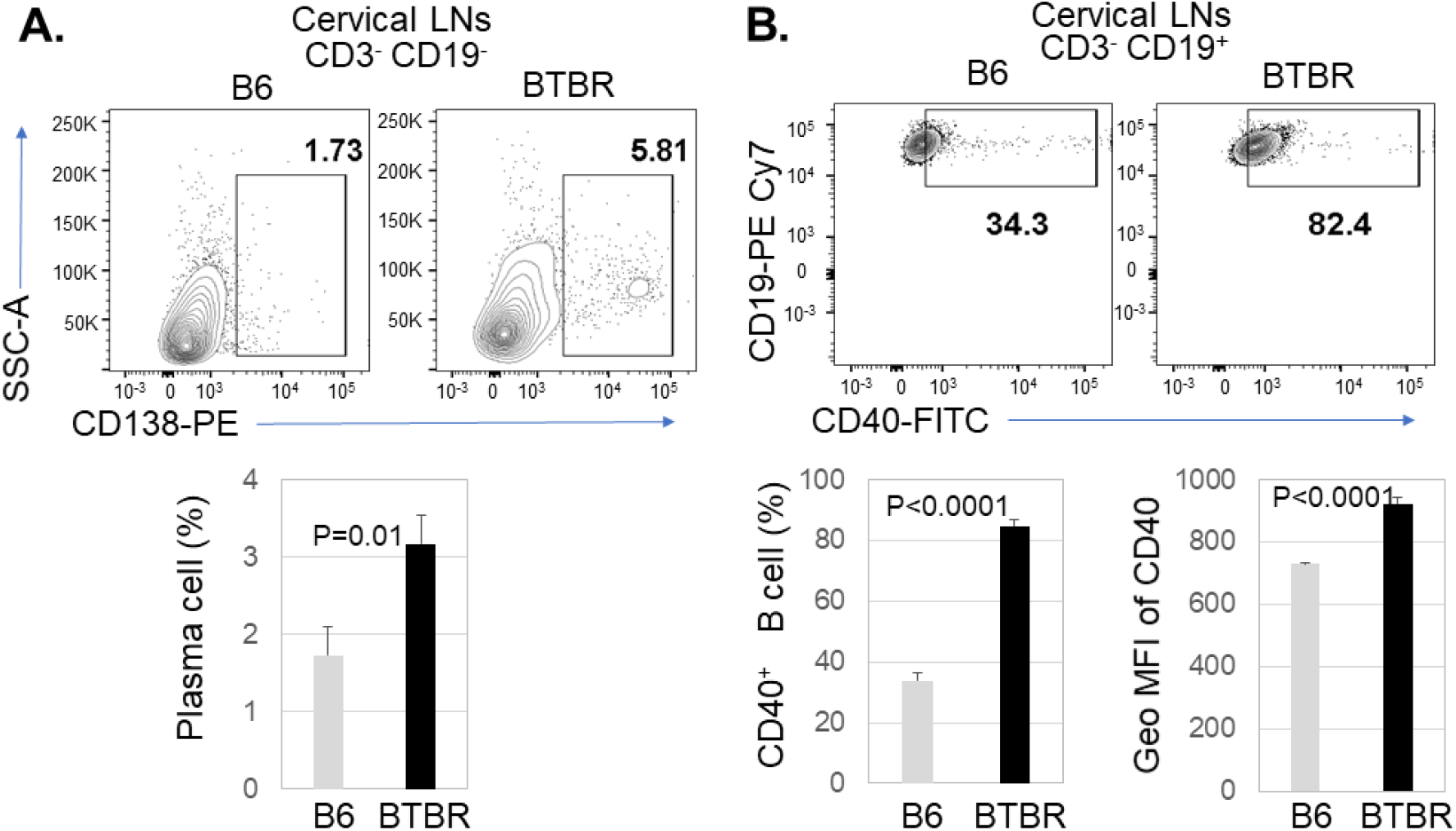
Plasma cells and activated B cells are increased in BTBR mice. Representative flow cytometric analysis and frequency of CD138^+^ plasma cells (A) and CD40 expression on B cells (B) in the cervical lymph nodes of B6 and BTBR mice. The geometric mean fluorescent intensity (GeoMFI) of CD40 for expression level on B cells in the cervical lymph nodes of B6 and BTBR mice is also shown (B). Data are representative of three independent experiments with 4-6 pairs of mice in each experiment. The *p* values were determined by unpaired two-tailed Student’s t test, and *p*<0.05 is considered as significantly different. Error bar indicates mean ± SEM.

### Serum IgG autoAb levels

BTBR mice have increased levels of total IgG and autoAbs to brain homogenate [40,41,44], but Ag specificities had not been assayed. Since BTBR spleens and cervical lymph nodes had higher number of plasma cells, we further assayed autoAbs for some specificities of Ags suggested to affect brain functions. Whole brain homogenates from SCID mice were used as Ags to select and detect IgG Abs to the captured Ags. The Abs are considered autoAbs since they are constitutively produced; the sera autoAbs levels are presented as optical density (OD) values.

Aquaporin (AQP)-4 is a major membrane water channel in the central nervous system. The IgG autoAb levels to AQP-4 were significantly higher in the sera of BTBR mice compared to B6 mice (Fig: 5A, P<0.0001). Similarly, the autoAb IgG levels to N-methyl-D-aspartate Receptor (NMDAR), Contactin Associated Protein 2 (CASPR2), a membrane protein complexed with the neuronal potassium channel (VGKC), and Pentraxin2/SAP were assayed. The ELISA data indicates that the levels of IgG autoAbs to NMDR, CASPR2 and pentraxin2/SAP were also significantly higher in BTBR sera compared to B6 sera (Fig: 5B-D, P<0.0001).

**Figure 5.**
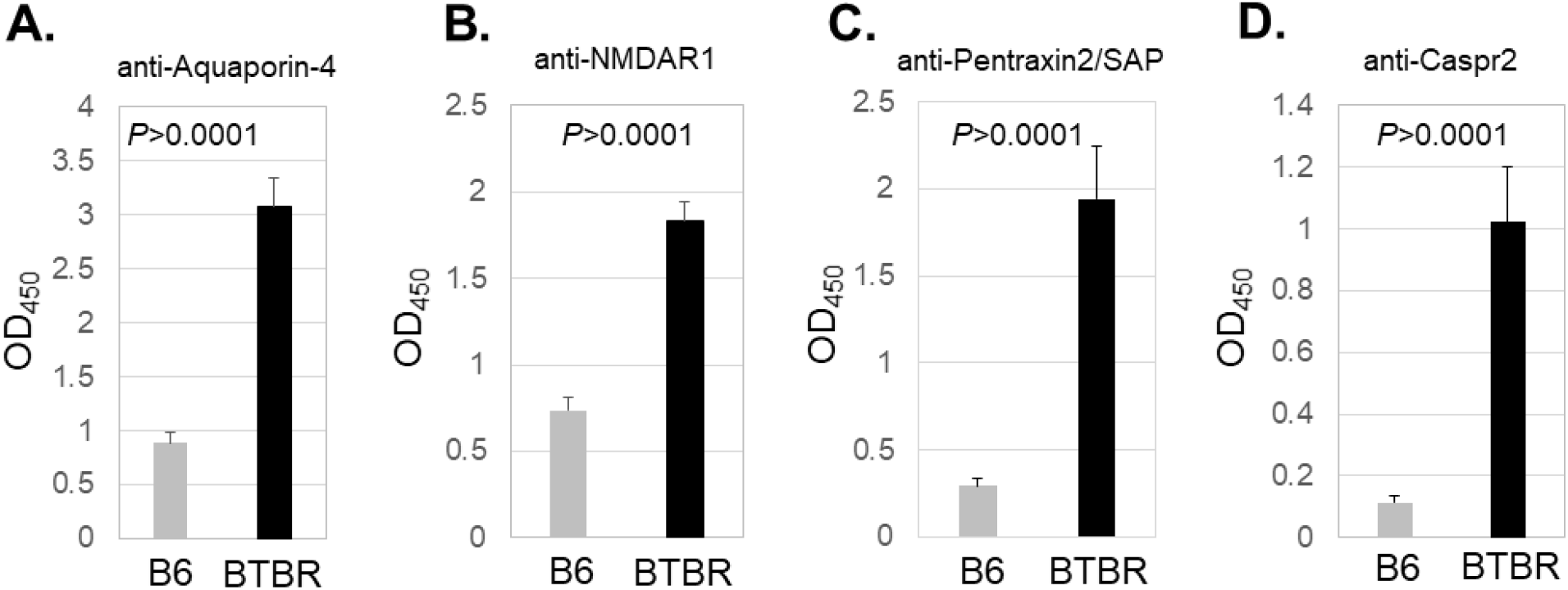
Increased level of serum IgG autoAbs to brain associated antigens in BTBR compared to B6 mice. ELISA of BTBR and B6 serum IgG Abs to Aquaporin-4 (A), NMDAR1 (B) Pentraxin (C) and Caspr2 (D). A 96-well plate was coated with 1 μg rabbit polyclonal Ab to Aquaporin-4, NMDAR1, Pentraxin and Caspr2 (D), blocked and used to capture Ag from 10 μg SCID brain homogenate for assessment of IgG anti-Aquaporin-4, anti-NMDAR1, anti-Pentraxin2/SAP and anti-Caspr2 in BTBR or B6 sera, which was diluted 1:10, and detected with HRP conjugated goat anti-mouse IgG Fc. Data are representative of four independent experiments with 6 pairs of mice in each experiment. The *p* values were determined by unpaired two-tailed Student’s t test, and *p*<0.05 is considered as significantly different. Error bar indicates mean ± SEM.

### AutoAb IgG to double-stranded (ds) DNA

The increased levels of antinuclear Abs have been described in children with autism. Analysis of serum anti-dsDNA Ab levels is a routine clinical assay for some autoimmune diseases. Thus, BTBR vs B6 sera were measured for IgG autoAbs to dsDNA. The ELISA data demonstrated that the IgG anti-dsDNA levels were higher in BTBR sera compared to B6 sera (Fig: 6, P<0.0001).

**Figure 6.**
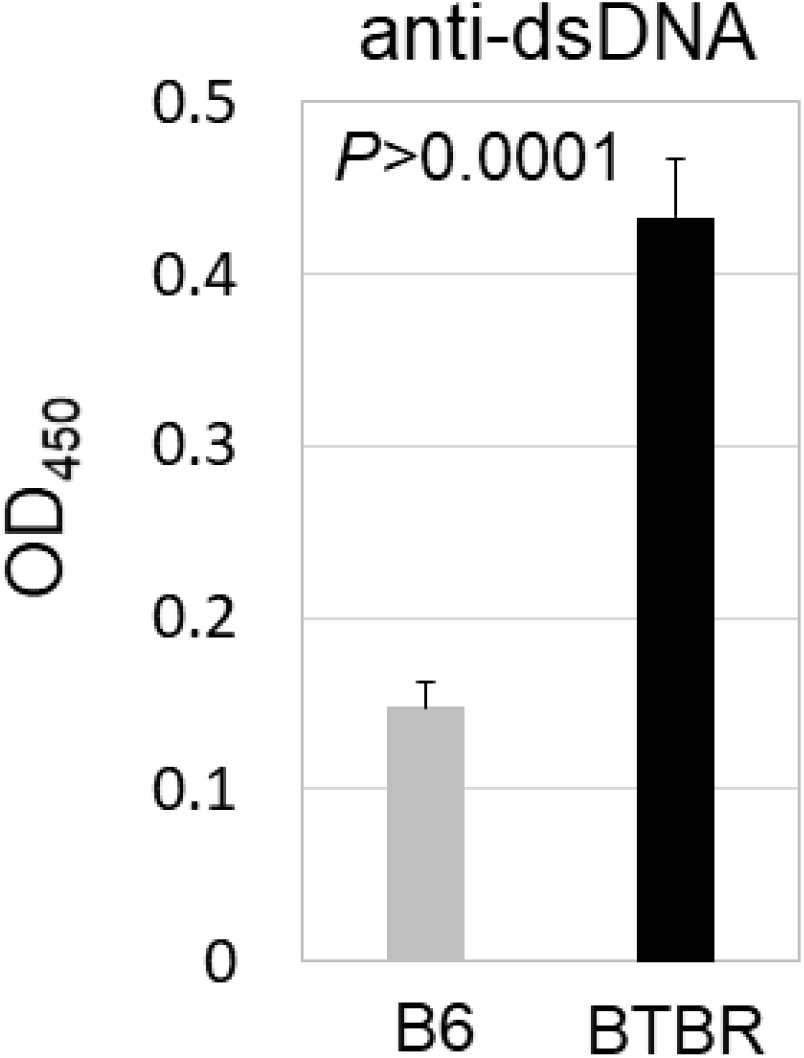
Increased serum anti-dsDNA levels in BTBR than B6 mice with ELISA analyses. UV-treated 96-well plates were used to coat with poly dA-dT to capture the anti-dsDNA in BTBR or B6 sera, which was diluted 1:100, and detected with HRP conjugated goat anti-mouse IgG Fc. Data are representative of four independent experiments with 6 pairs of mice in each experiment. The *p* values were determined by unpaired two-tailed Student’s t test, and *p*<0.05 is considered as significantly different. Error bar indicates mean ± SEM.

### T regulatory (Treg) cells in the meninges and lymphoid organs

Since the activation status of T cells and B cells, abundance of plasma cells and autoAb level are higher in BTBR mice, it was important to assess if this could be accounted for by a Treg deficiency. We investigated the Treg cell population in the meninges, cervical lymph nodes and spleen based on the expression of CD25 and FoxP3 in the CD4^+^ T cell population. Flow cytometric analysis revealed the level of Treg cells in the meninges (Fig: 7A, P=0.001) and cervical lymph nodes (Fig: 7B, P=0.0005) were significantly lower in BTBR mice compared to B6 mice. However, the Treg frequency in the spleen was increased (Fig: 7C, P=0.0006).

**Figure 7.**
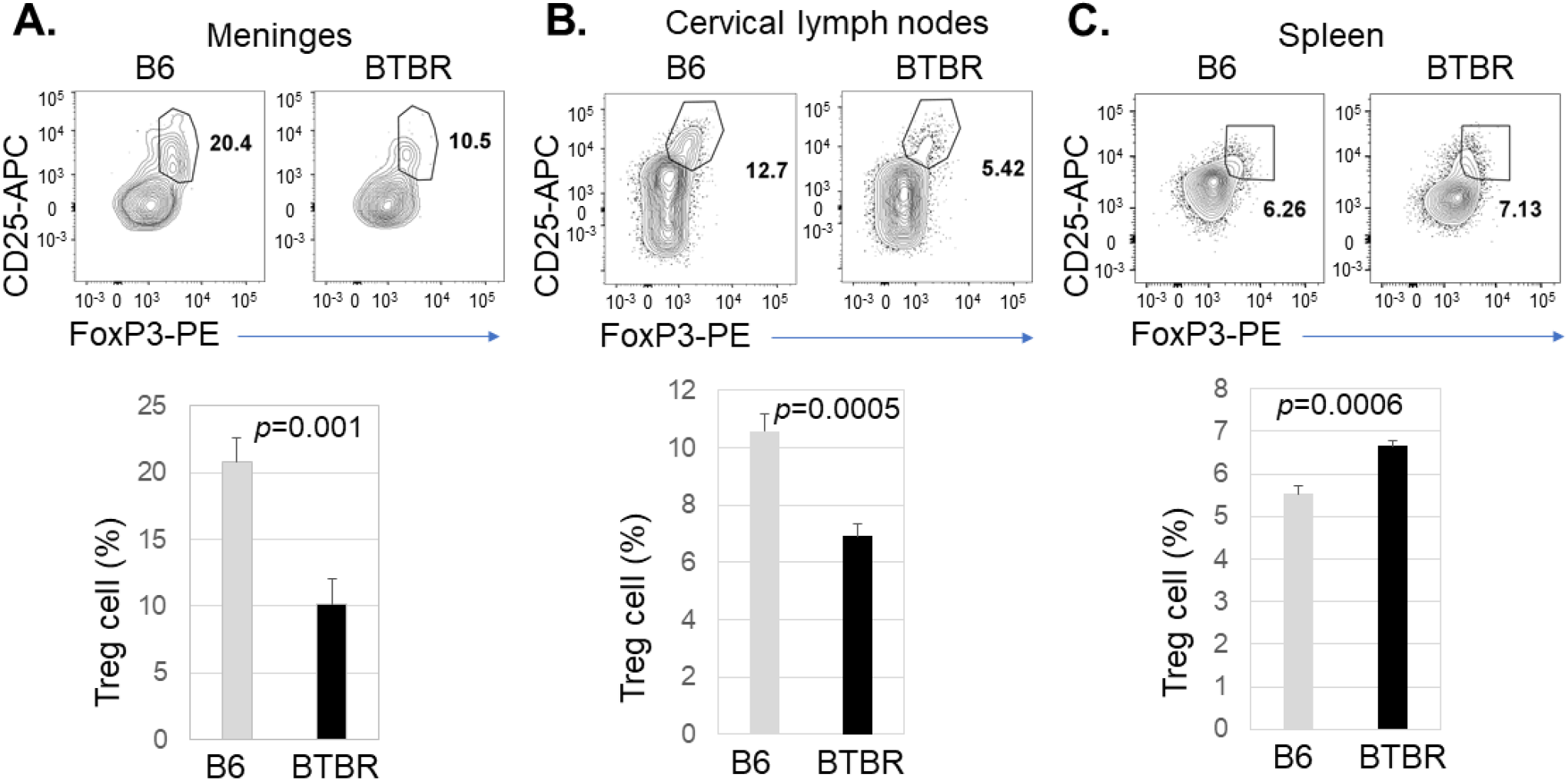
Decreased frequency of Treg cells in the meninges and cervical lymph nodes of BTBR mice. Representative flow cytometric analysis of CD25^+^FoxP3^+^ Treg cells in the meninges (A), cervical lymph nodes (B) and spleen (C) of B6 and BTBR mice. Frequencies of CD25^+^FoxP3^+^ T regulatory cells (Treg) cells in the meninges (A), cervical lymph nodes (B) and spleen (C) of B6 and BTBR mice are shown. Treg cell population were identified based on CD25^+^FoxP3^+^ cell in CD3^+^CD4^+^ population. To observe the Treg cell population, CD4^+^ cells were analyzed for the expression of CD25 and FoxP3 and gated on the double positive population. Data are representative of three or four independent experiments with 6 pairs of mice in each experiment. P values determined by unpaired two-tailed Student’s t test are indicated and p<0.05 is considered as significantly different. Error bars indicate mean ± SEM.

### Cytokine production by Treg cell

Since there were decreased levels of Treg cells in meninges and cervical lymph nodes but increased in the spleen, Treg function based on IL-10 expression was also necessary. To investigate Treg function, cells were stimulated *in vitro* and assayed for production of the immunosuppressive cytokine IL-10. The frequency of Treg cells producing IL-10 and IL-10 expression level were significantly higher in the cells from meninges (Fig: 8A, P<0.0001), cervical lymph nodes (Fig: 8B, P<0.0001 & P=0.0003) and spleen (Fig: 8C, P<0.006 & P=0.032) of BTBR mice compared to B6 mice. Although the spleen showed higher frequency of Treg based on percentage of CD4^+^ T cells expressing CD25 and FoxP3, after stimulation in vitro the splenic cultures had fewer CD4^+^ cells expressing IL-10 and their IL-10 expression was lower (Fig.9).

**Figure 8.**
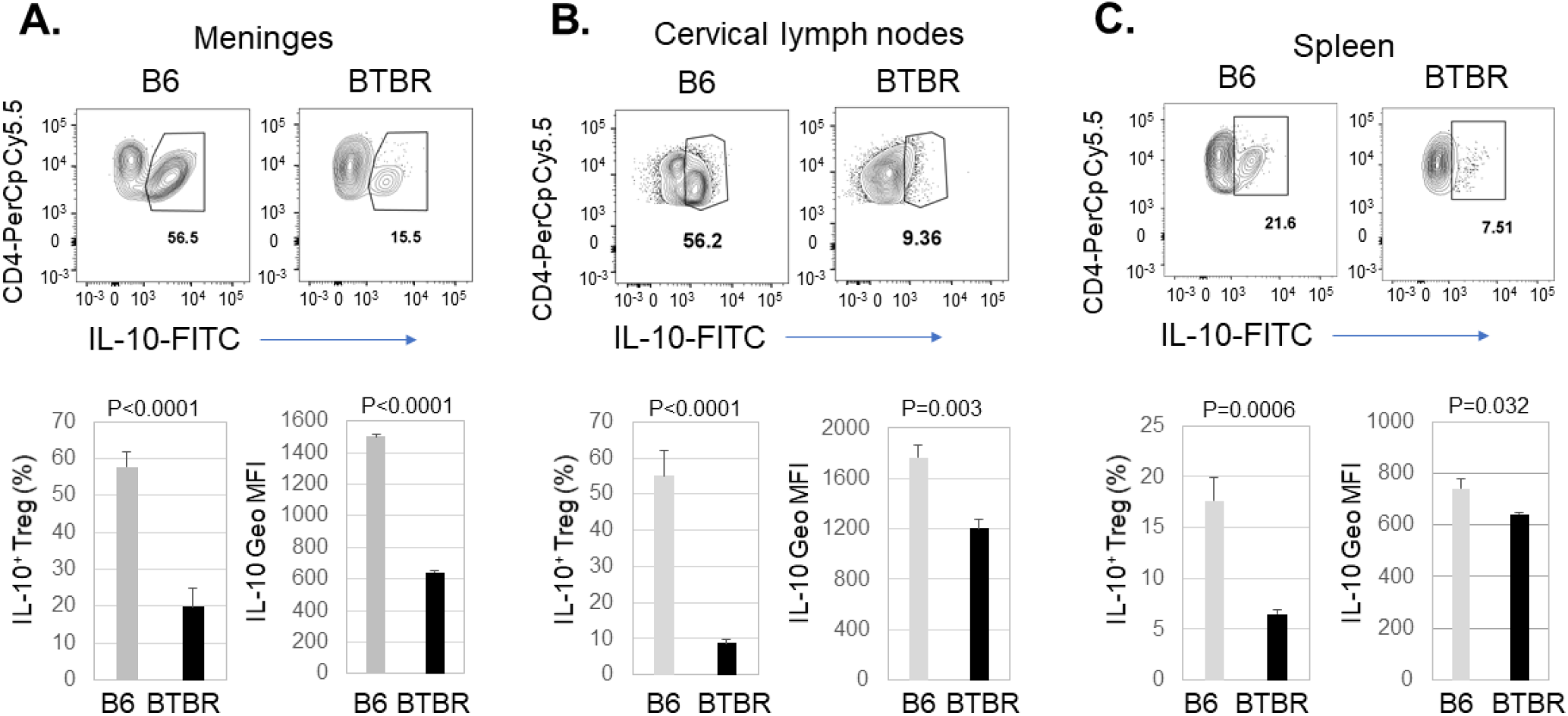
Decreased production of IL-10 by Treg cells in BTBR mice. Representative flow cytometric analysis of IL-10 production by CD25^+^FoxP3^+^ T regulatory cells (Treg) cells in the meninges (A), cervical lymph nodes (B) and spleen (C) of B6 and BTBR mice. Frequencies of IL-10^+^ Treg cells and geometric mean fluorescent intensity (Geo MFI) of IL-10 in Treg cells from meninges (A), cervical lymph nodes (B) and spleen (C) of B6 and BTBR mice are shown. Cells were stimulated in vitro with Cell activation cocktail, stained with cell surface marker and then stained with intra-cellular cytokine IL-10. Data are representative of three independent experiments with 4-6 pairs of mice in each experiment. P values determined by unpaired twotailed Student’s t test are indicated and p<0.05 is considered as significantly different. Error bars indicate mean ± SEM.

**Figure 9.**
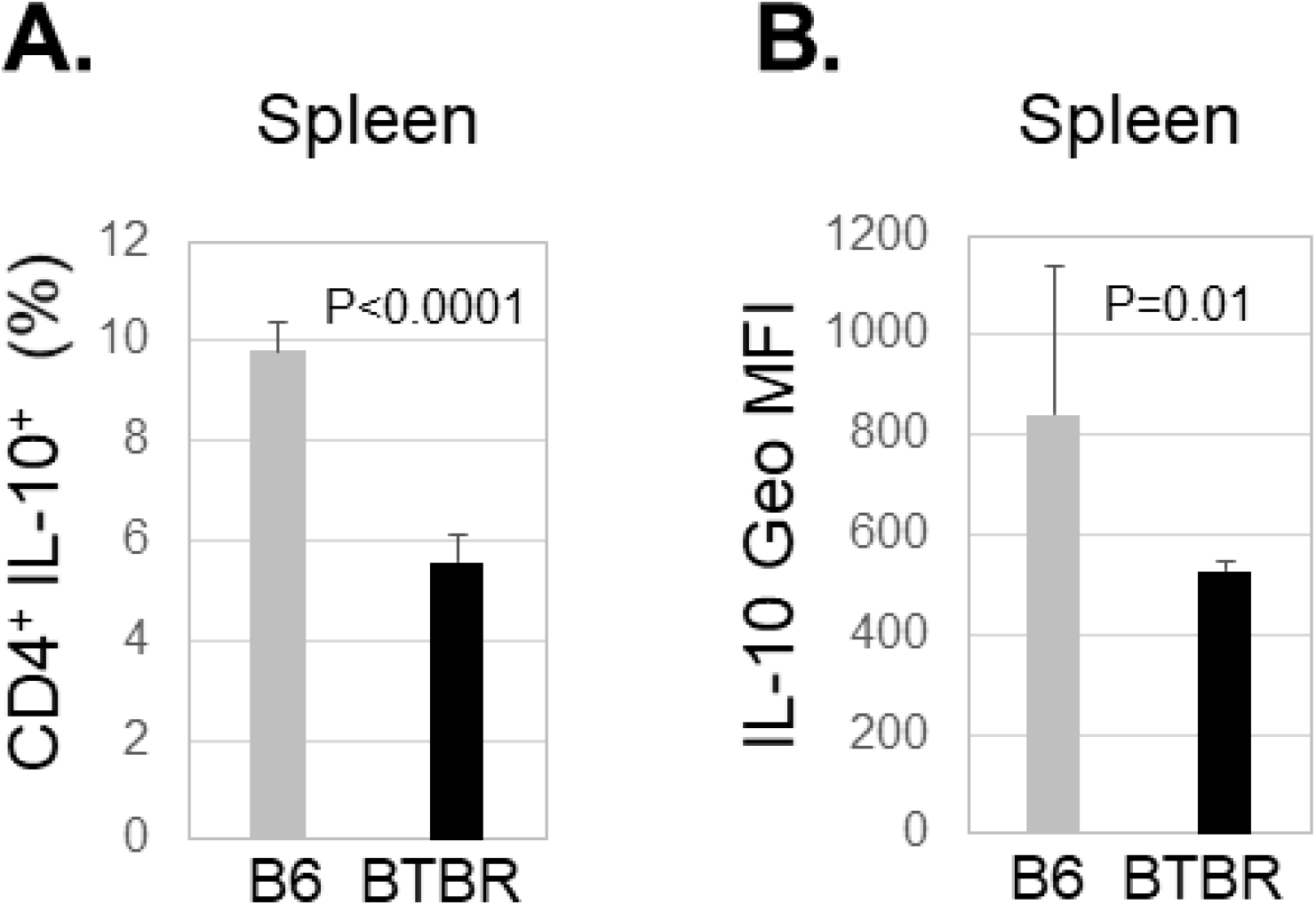
Decreased production of IL-10 by CD4^+^ cells in BTBR mice. Frequencies of CD4^+^ IL-10^+^ cells (A) and geometric mean fluorescent intensity (Geo MFI) of IL-10 (B) in CD4^+^ cells from spleen of B6 and BTBR mice. Cells were stimulated in vitro with Cell activation cocktail, stained with cell surface marker and then stained with intra-cellular cytokine IL-10. Data are representative of three independent experiments with 4-6 pairs of mice in each experiment. P values determined by unpaired two-tailed Student’s t test are indicated and p<0.05 is considered as significantly different. Error bars indicate mean ± SEM.

## Discussion

This study examined the meningeal immunoprofile (immune cell types, cell activation status, and cell function) of the BTBR mouse model of ASD and autoAbs to Ags that might have an influence on neurodevelopment and neurofunctions. Mechanisms involved in autoimmunity also were considered. BTBR mice serve as a widely recognized mouse model as they display unusual behavior including lack of social interaction and restricted repetitive actions that bear a resemblance to ASD [38,46]. Extensive modifications of immune functionality have been observed in individuals with ASD, such as neuroinflammation, higher proinflammatory cytokines in the brain and peripheral blood, brain Ag-specific autoAbs and altered immune profiles [50]. Moreover, these dysfunctional immune responses relate to increased weakening of behaviors and deficits in social relations and communication, suggesting that the immune system plays a key role in the development of ASD. [51,52]. AutoAbs to neuronal Ags have been reported in numerous autoimmune neuronal disorders including ASD [53,54].

Immune cells such as T cells, B cells, innate lymphoid cells (ILCs), macrophages, dendritic cells, mast cells, and neutrophils can circulate through the meningeal lymphatic system under usual and pathological situations. Their activation, functions and signals are important for brain homeostatic conditions and animal behavior [10,55]. Autoimmune T cells and B cells in meninges can lead to neuronal inflammation and motor or intellectual dysfunction [56,57]. A recent study proposed that meningeal immunity might control the neuroinflammatory response in autoimmune disease including multiple sclerosis (MS) [12]. Autoreactive CD4^+^ T cells can enter the CNS through the meningeal blood vessels, where they are restimulated by Ag presenting cells (APC), which can promote the development of neuroinflammation by producing inflammatory cytokines [58]. An inflammatory condition can induce a tertiary lymphoid structure in meningeal space, which can attract and activate T cells, B cells and APCs leading to pathogenic condition in neuronal autoimmune diseases [59].

An increased proportion of T cell and B cell populations are present in the meninges of BTBR mice compared to B6 mice. Moreover, T cells in BTBR meninges are activated, indicated by higher levels of CD25 expression. Higher level of MHCII molecule expression on B cells and myeloid cells also indicated greater activation in BTBR meninges. BTBR mice have higher levels of serum IgG and autoAbs to brain associated Ags, which bound to different brain regions and fractions of brains and neuronal cell lines such as MN9D and CATH.a [40, 44]. Activated B cell (CD19^+^CD40^+^) and CD138^+^ plasma cell population may contribute to the autoimmune condition in BTBR mice.

BTBR mice had higher serum IgG levels to AQP-4, NMDAR1, Caspr2, and pentraxin. Anti-AQP-4 Abs have been linked to an inflammatory Neuromyelitis Optica Spectrum Disorder (NMOSD) that specifically disrupts optic nerves and spinal cord [60]. AutoAb to NMDAR is known to cause dysfunctional glutamate neurotransmission in the brain that manifests as psychiatric symptoms [61]. Brimberg et al. [62] reported an Ab binding to Contactin Associated Protein 2 (CASPR2), a membrane protein complexed with the neuronal potassium channel, is abundant in a mother of ASD child who displayed abnormal neuronal development as well as weakening social interaction, flexible learning and monotonous behavior. AutoAbs to pentraxins have been observed in systemic lupus erythematosus (SLE) and many other autoimmune diseases [63]. SAP can suppress development of experimental autoimmune encephalomyelitis [64]. Thus, BTBR Abs have multiple specificities that may be responsible for the ASD-like behaviors. Brain-reactive Abs are higher in mothers of ASD individuals and are suggested to connect with autoimmunity [65]. The BTBR mice also have a high ratio of antinuclear immunoglobulins. Anti-dsDNA or anti-nucleosome Abs are present in 34 and 47% of individuals with ASD [66] and SLE patients. [67].

The low functionality of Treg cells and higher number of splenic Tfh cells [47] in the BTBR mice allow the differentiation of more B cells to become plasma cells generating the autoAbs interfering with brain functions. Treg cells as a primary mediator of peripheral immune tolerance, protecting from autoimmune diseases, and restricting chronic inflammatory diseases [68] were dysfunctional in the BTBR mice. FOXP3 expressing Treg cells are important for immune homeostasis and a reduction of this population is connected to up-regulation of many cell types leading to inflammatory neuronal disorder [36,40]. Anti-inflammatory cytokine IL-10 produced by Treg cells plays an important role in suppressing the immune response [69]. BTBR mice having fewer Treg cells and less IL-10 in the meninges and cervical lymph nodes allows more inflammation. With more neuroinflammation, there may be enhanced leakage of brain Ags causing more damage associated molecular patterns (DAMPS) to stimulate microglia and other myeloid cells to release cytokines and chemokines attracting more T and B cells to the meninges or even plasma cells into the brain parenchyma. A similar outcome may occur in individuals with ASD and in experimental mouse model of ASD [4,46]. Although BTBR mice have an elevated number of Treg cells in spleen, their production of IL-10 was low like that of the Treg cells in BTBR meninges and cervical lymph nodes. Bone marrow derived macrophages from BTBR mice also produce a lower level of IL-10 than B6 mice [49]. Moreover, IL-10 suppresses expression of MHC-II molecule and co-stimulatory molecules on antigen presenting cells (APCs) such as DCs and macrophages to control the proliferation of antigen-specific CD4^+^ T cells [70]. Overall, lower level of Treg cells and their decreased functional activity is unable to prevent the autoimmune-like phenotype of BTBR mice and could possibly contribute to the autism-like behavior.

Male BCF1 mice showed higher anti-brain Ab sera levels than those of B6 mice, however, they had lower levels than the BTBR mice, had less IgG in the brain and no significance differences from B6 mice were observed in social interaction and behavioral index [43]. Although adult BTBR progeny that developed in B6 dams had similar adult levels of serum anti-brain Abs and IgG in the brain as BTBR mice, they had improved behavior [44]. The influence of neonatal experience in the maternal environment is important, and apparently anti-brain Abs alone is only partially responsible for the aberrant behaviors. However, serum BTBR IgG given to B6 dams on gestational days 13-18 was able to significantly lower the normal behavior of the adult offspring [44]. The pathogenesis of autoimmune diseases is connected to genetic disposition and epigenetic modifications. Damage in central and peripheral immune tolerance results in autoimmune diseases including systemic lupus erythematosus, type 1 diabetes, rheumatoid arthritis, and primary biliary cirrhosis [71]. Genes such as *AIRE, Foxp3, CTLA4* and *FAS* are associated with loss of immune tolerance in autoimmunity [72]. Additionally, epigenetics changes such as DNA methylation, histone modification and alteration in microRNAs expression are possibly accountable for the failure of immune tolerance in autoimmune disorders [72]. Since the maternal environment plays a significant role in the early development affecting later abnormal behaviors, investigating maternal nuclear and mitochondrial inheritance genes as well as early maternal epigenetic changes during fetal development could potentially decipher the mechanism of autism development. The reduced levels of Treg cells and IL-10 and their effects on the T and B cell populations are likely implicated in the development of autism-like behavior with production of autoAbs; however, other immune cell types and brain neurotrophic factors might be involved. The lower production of IL-10 and higher levels of inflammatory cytokines such as IL-1β, IL-18 and IL-33, which are higher in BTBR mice [43] may account for the neurodevelopment disorder.

## Materials and methods

### Animals

Two-to three-month-old C57BL/6J (B6) and BTBR *T*^+^ *Itpr3^tf^*/J (BTBR) mice were obtained from Jackson Laboratories (Bar Harbor, ME, USA). Both genders were used for all experiments. Findings were combined across sex because of no differences between males and females were obtained. Mouse colonies were maintained in the AAALAC-approved Wadsworth Center Animal Facility under ideal conditions of humidity and temperature in a 12h light-dark cycle (7AM-7PM). All procedures were approved by Wadsworth Center’s IACUC.

### Cell and serum preparation

Spleens, cervical lymph nodes and brain meninges were harvested after perfusion with cold phosphate buffered saline (PBS). Single cell suspensions were prepared from spleens and cervical lymph nodes by pressing through the cell strainer (Fisherband, Sterile Cell strainer, 100 uM Cat No: 22363549, Fisher Scientific). Red blood cells (RBC) were lysed with RBC lysis buffer (Cat; 00-4333-57, eBioscience/Invitrogen) before staining.

Cells from the meninges were collected according to previously published protocol with few modifications [73]. Briefly, using sharp forceps the meninges were collected under dissecting microscope and transfer into Petri dish containing RPMI supplemented with 25mM HEPES. Meninges were digested with Collagenase D (Cat:11088858001, Roche, Germany) and DNase I (Cat:10104159001, Roche, Germany) for 1 hr at 37°C on rotator. The digested meninges solution passed through the nylon mesh strainer in a 50 mL conical tube with the help of a 5 ml syringe plunger. The strainer was washed with an additional 2 to 3 mL of RPMI/HEPES media to pass the cells through the strainer. To get the suitable cell numbers, meninges from 3-4 animals were pooled together. Cells were counted with an automated cell counter (Countess II FL, Invitrogen) after staining with viability dye, and 1 x 10^6^ cells were used for staining.

Blood samples were collected into centrifuge tubes and kept at room temperature for 30-60 min. Sera were separated by centrifugation at 14,000 ×*g* for 10 min. Aliquoted serum was stored at −80°C until use.

### *In vitro* stimulation for cytokine production

For *in vitro* cytokine production assay, cells from spleen, cervical lymph nodes and meninges were placed in 96-well plates, at a concentration of 1 × 10^6^ cells per 100 μl in RPMI medium supplemented with 10%FBS and stimulated for overnight with Cell activation cocktail (Cat:423301, Biolegend) at 37°C incubator. Brefeldin A solution (Cat: 420601, Biolegend) and Monensin solution (Cat: 420701, Biolegend) were added to the cells for the final 3 hrs. Cells were collected, stained for surface markers and intracellularly for cytokines, and analyzed by flow cytometry.

### Surface and intracellular/intranuclear staining

Cells from the spleen, cervical lymph nodes and meninges were transferred into FACS tube (100 μl, 1X10^6^ cell), blocked with FC blocker (anti-CD16/32) and stained with surface antibodies. Cells were then washed, resuspended with FACS buffer and acquired by Flow Cytometer. For intracellular cytokine staining, cells were stained for surface markers in the presence of Brefeldin A solution (Cat: 420601, Biolegend) and Monensin solution (Cat: 420701, Biolegend). After washing cells were fixed and permeabilized with BD Cytofix/Cytoperm solution (Cat: 51-2090KZ, BD Bioscience), and washed with BD Perm/Wash solution (Cat: 51-2091KZ, BD Bioscience). After resuspension in Perm/Wash solution, cells were stained for intracellular cytokine, washed, resuspended in FACS buffer and acquired by FACSCanto flow cytometer. For intranuclear staining (FoxP3), cells were stained with surface antibody and then fixed and permeabilized with fixation/permeabilization buffer (Fixation/Permeabilization buffer set, Cat: 00-5123-43, eBioscience). Intranuclear staining and washing were performed in the presence of permeabilization buffer (Permeabilization buffer, Cat:00-8333-56, eBioscience). After final washing cells were resuspended in FACS buffer and data were acquired by FACSCanto flow cytometer (BD Biosciences)

### Flow cytometry

The cells were stained for surface and intracellular/intranuclear, and analysis by FACS Canto flow cytometer (BD Biosciences). The following fluorochrome conjugated Abs: PerCP Cy5.5-CD45, APC Cy7-CD45, PerCP Cy5.5-CD4, PE-CD4, FITC-CD4, FITC-CD3, APC-CD3e, PerCP-CD3e, APC Cy7-CD3, PE-CD11b,PE Cy7-CD11b, FITC-MHC-II, PerCP Cy5.5-MHCII, PE-CD19, FITC-CD19, PE Cy7-CD19, APC-CD25, PE Cy7-CD25, PerCp Cy5.5-CD25, APC-CD138, FITC-CD40, PE-FoxP3, FITC-IL10 and anti-CD16/32 (Fc block) were purchased from BD Pharmingen (San Diego, CA), Biolegend (San Diego, CA), or eBiosciences (Thermo Fisher Scientific, MA, USA). Frequencies and numbers of populations in the meninges, lymph nodes and spleens of B6 and BTBR mice were gated based on FSC-A and SSC-A, followed by gating in single cells and finally gated on CD45^+^ cells. Acquired FCS files were analyzed by Flow Jo-V10.

### ELISA for IgG anti-dsDNA in mouse serum

The level of total anti-dsDNA in serum was determined with a sandwich ELISA. Costar assay plate (96-well, flat bottom, High binding. Corning incorporated, Ref: 3369) were U-V treated and coated with poly dA-dT (Sigma D4522, Lot: SLCD4909, 7.5 μg/ml) and HRP-conjugated goat anti-mouse IgG Fc (dilution 1:10000, Cat: 115035008, Jackson immune research) was used as the detection Ab; TMB (3,3’,5,5’-tetramethylbenzidine) was used as substrate (Sigma). Plates were washed with an automated plate washer (BioTek ELx405Select CW, Winooski, VT) and read the absorbance at OD_450_ with an ELISA plate reader (BioTek EL808). Sera were diluted 1:100 with 10% FBS in PBS.

### ELISA for IgG autoAbs

Rabbit polyclonal anti-AQP4 (SAB5200112-100UL, Lot:PA187526, Millipore Sigma), rabbit polyclonal anti-Pentraxin2 or SAP (R&D systems, Catalog # AF2558), rabbit polyclonal anti-Caspr2/CNTNAP2 (Cat: ab218048, abcam), and rabbit monoclonal anti-NMDAR1 (Clone # 54.1, Invitrogen, Catalog # 32-0500, Lot: WA314671) were used to coat 96-well plate (0.5 μg/well) for capturing Ags known to be expressed in brains. The plates were incubated for 2 hr at room temperature. After 3X washing, the plates were blocked with 2.5% BSA-PBST for overnight at 4°C. After incubation and washing, SCID mouse whole brain lysate (10 ug/100ul/well) was added into the wells and incubated for 2 hr at room temperature. After 3X washing, sera from B6 or BTBR mice (dilution 1/10 or 1/100) were added into the wells and incubated for 2 hr at room temperature. After 6X washing, HRP-goat anti-mouse IgG Ab (dilution 1:10000, Cat: 115035008, Jackson immune research) was added and incubated for another 2 hr at room temperature. The plates were washed 6X, and the TMB (3,3’,5,5’-tetramethylbenzidine, Sigma) was used as a substrate solution for color development. Absorbance was measured at 450 nm by ELISA analyzer (BioTek EL808).

### Statistical analysis

Data are presented as mean ± SEM, unpaired two-tailed Student’s t test were used to determine p values and *p*<0.05 measured as statistically significant difference.

## Acknowledgements

We acknowledge Dr. Yunyi Yao and the Wadsworth Center animal facility staff for their assistance for the maintenance of the mice.

